# Fitness costs of parasites explain multiple life history tradeoffs in a wild mammal

**DOI:** 10.1101/683094

**Authors:** Gregory F Albery, Alison Morris, Sean Morris, Fiona Kenyon, Daniel H Nussey, Josephine M Pemberton

**Affiliations:** University of Edinburgh; Moredun Research Institute

## Abstract

Reproduction in wild animals can divert limited resources away from immune defence, resulting in increased parasite burdens. A longstanding prediction of life history theory states that these parasites can harm the reproductive individual, reducing its subsequent fitness and producing reproduction-fitness tradeoffs. Here, we examined associations among reproductive allocation, immunity, parasitism, and subsequent fitness in a wild population of individually identified red deer (*Cervus elaphus*). Using path analysis, we investigated whether costs of lactation for downstream survival and fecundity were mediated by changes in strongyle nematode count and mucosal antibody levels. Lactating females exhibited increased parasite counts, which were in turn associated with substantially decreased fitness in the following year in terms of overwinter survival, fecundity, subsequent calf weight, and parturition date. This study offers observational evidence for parasite regulation of multiple life history tradeoffs, supporting the role of parasites as an important mediating factor in wild mammal populations.

## Introduction

A fundamental tenet of life history theory states that reproduction should be costly for future fitness (Williams, 1966; Stearns, 1989). While evidence for such trade-offs is widespread, the mechanisms behind them remain poorly understood. One hypothesised mechanism is that reproductive costs act through parasites, where increased reproductive allocation diverts limited resources away from immune defence, resulting in increased parasite burdens, which reduce subsequent fitness (Sheldon and Verhulst, 1996; Harshman and Zera, 2007). Parasite mediation of life history tradeoffs involves two necessary components: that reproduction increases parasitism, and that these parasites cause harm or require resources to combat them, reducing subsequent fitness. There is abundant evidence for each component of this theory across a range of taxa: firstly, life history investment is often associated with weaker or altered immune allocation (Neggazi *et al.*, 2016; Rödel *et al.*, 2016; Krams *et al.*, 2017) or with increased parasitism (Festa-Bianchet, 1989; Cizauskas *et al.*, 2015; Debeffe *et al.*, 2016). Secondly, increased parasitism is often associated with decreased subsequent probability of survival (Coltman *et al.*, 1999; Leivesley *et al.*, 2019) or reproduction (Albon *et al.*, 2002; Vandegrift *et al.*, 2008; Hughes *et al.*, 2009). Despite evidence for one or other of these processes in isolation, reproduction-associated increases in parasitism have rarely been linked to downstream fitness consequences in the same study to provide full support for parasite mediation of life history tradeoffs.

While it is true that reproduction, immunity, and parasites all compete for host resources, mechanisms governing life history tradeoffs are hypothesised to occur in a temporal sequence rather than occurring simultaneously (Figure 1). First, reproduction diverts resources away from immunity, reducing immune allocation (Sheldon and Verhulst, 1996). Resultant weaker immunity, plus potentially increased exposure associated with altered behaviour of reproductive individuals, can then result in higher parasite burden (Knowles *et al.*, 2009; Albery *et al.*, 2020). Finally, subsequent fitness is reduced by damage from parasites (Harshman and Zera, 2007; Graham *et al.*, 2011). This combination of mechanisms comprises an indirect cost of reproduction acting through parasites. Additional (direct) costs of reproduction can simultaneously act through other mechanisms such as reduced condition, hormonal or phenological regulation, or damage caused by oxidative stress (Stjernman *et al.*, 2004; Harshman and Zera, 2007; Speakman, 2008; Figure 1). This causal sequence is important, because parasites’ observed relationship with life history traits can depend on whether a preceding, contemporary, or subsequent trait is chosen to examine.

**Figure 1:**
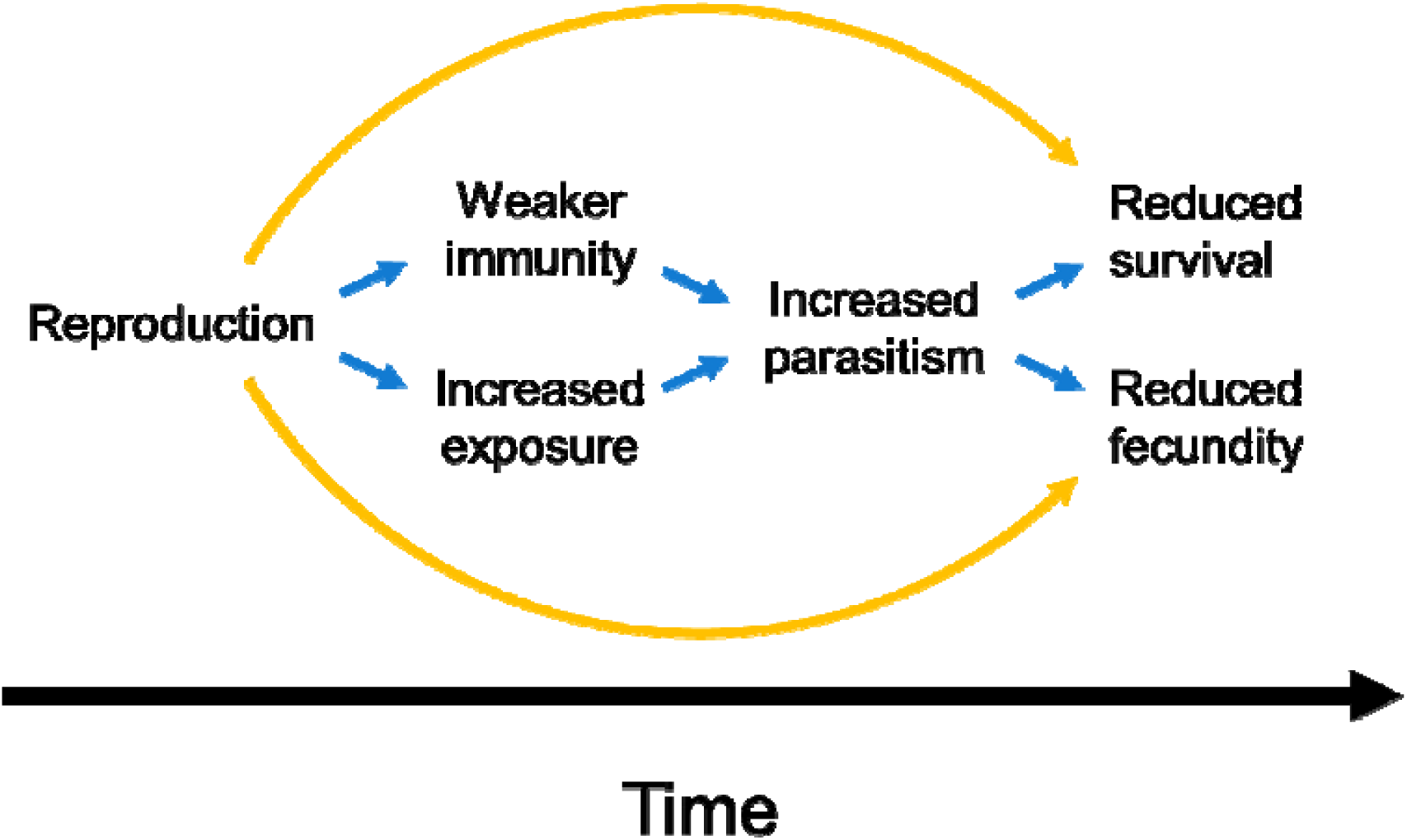
The hypothesised mechanism for parasite-dependent mediation of life history tradeoffs. Blue (interior) arrows denote indirect, parasite-mediated fitness costs of reproduction, while the orange (exterior) arrows denote direct costs through resource allocation, hormonal regulation, or similar mechanisms.

Many studies examining reproduction-immunity-parasitism interrelationships have been carried out in birds, often using experiments in which reproductive effort is artificially increased by manipulating clutch sizes (Knowles *et al.*, 2009). Such manipulations are not possible in many mammal species, and thus most of our knowledge of these trade-offs are based on observational studies. In observational contexts, or in concert with experiments, path analysis can be used to infer links between parasites and their fitness consequences (Pacejka *et al.*, 1998; Stjernman *et al.*, 2004; Brambilla *et al.*, 2015; Leivesley *et al.*, 2019). Notably, a recent analysis in a wild population of Soay sheep used path analysis to demonstrate observationally that reproduction reduced survival through increased parasite count and reduced body weight (Leivesley *et al.*, 2019), but without examining impacts on subsequent reproductive traits.

The wild red deer (*Cervus elaphus*) on the Isle of Rum provide a classic example of a life history tradeoff under natural conditions: female deer that invest in lactation have reduced future fecundity and survival probability compared to non-lactating females (Clutton-Brock *et al.*, 1989; Froy *et al.*, 2016). Females that give birth to a calf that dies within the first few months of its life have similar fitness outcomes to those that do not give birth, implying that gestation has a minimal cost relative to lactation (Clutton-Brock *et al.*, 1989). The deer are infected with several helminth parasites, with egg counts and mucosal antibody (IgA) levels measured via noninvasive collection of faecal samples (Albery *et al.*, 2018, 2020). A previous study demonstrated decreased mucosal IgA and increased parasite count in reproductive females (Albery *et al.*, 2020). Lactation (but not gestation) is associated with increased parasite counts, partially reflecting the cost of reproduction for fitness; however, investment in gestation is associated with decreased mucosal antibody levels (Albery *et al.*, 2020). These findings each demonstrate a cost of reproduction for fitness, parasitism, and immunity respectively in this population. However, we have yet to establish the degree to subsequent fitness costs of reproduction can be explained by changes in parasitism.

Here, we use path analysis to link reproduction-immunity-parasitism tradeoffs in the Isle of Rum red deer with survival and reproduction in the following year, investigating whether immunity and parasitism are capable of mediating life history tradeoffs, and attempting to separate immune and parasite mediation from direct effects of reproduction acting through alternative mechanisms. We expected that substantially increased parasite counts associated with lactation would be associated with decreased subsequent survival, fecundity, parturition date, and calf birth weight, so that parasites provide a mechanistic explanation for the costs of lactation seen in this system.

## Methods

### Study system, sampling, and labwork

The study population is situated in the north block of the Isle of Rum National Nature Reserve (57°N 6°20′W). The deer are entirely wild and unmanaged, and have been monitored continuously since the 1970s (see Clutton-Brock *et al.*, 1982 for an overview of the project). The life history data collected on the population provide high-resolution estimates of individuals’ dates of birth and death, reproduction, and familial relationships. The “deer year” begins on May 1^st^, and the deer give birth (“calving”) in May-June, having conceived in the previous autumn (Figure 2). Deer on Rum give birth to a single calf, and do not reproduce every year. During the calving season, we aim to capture and mark as many of the calves born as possible soon after birth, so that they can be monitored for the rest of their lives. Sex and capture weight (to the nearest 100g) are recorded. ∼20% of calves die within the first few weeks of life, and giving birth to a calf that dies within this period has little cost to the mother in terms of her survival and reproduction probability the following year (Clutton-Brock *et al.*, 1989). In contrast, if a calf survives into the winter, the mother has spent ∼6 months lactating to it, expending considerable resources in doing so, and this cost is associated with substantially decreased fecundity and survival probability the following year (Clutton-Brock *et al.*, 1989; Froy *et al.*, 2016).

**Figure 2:**
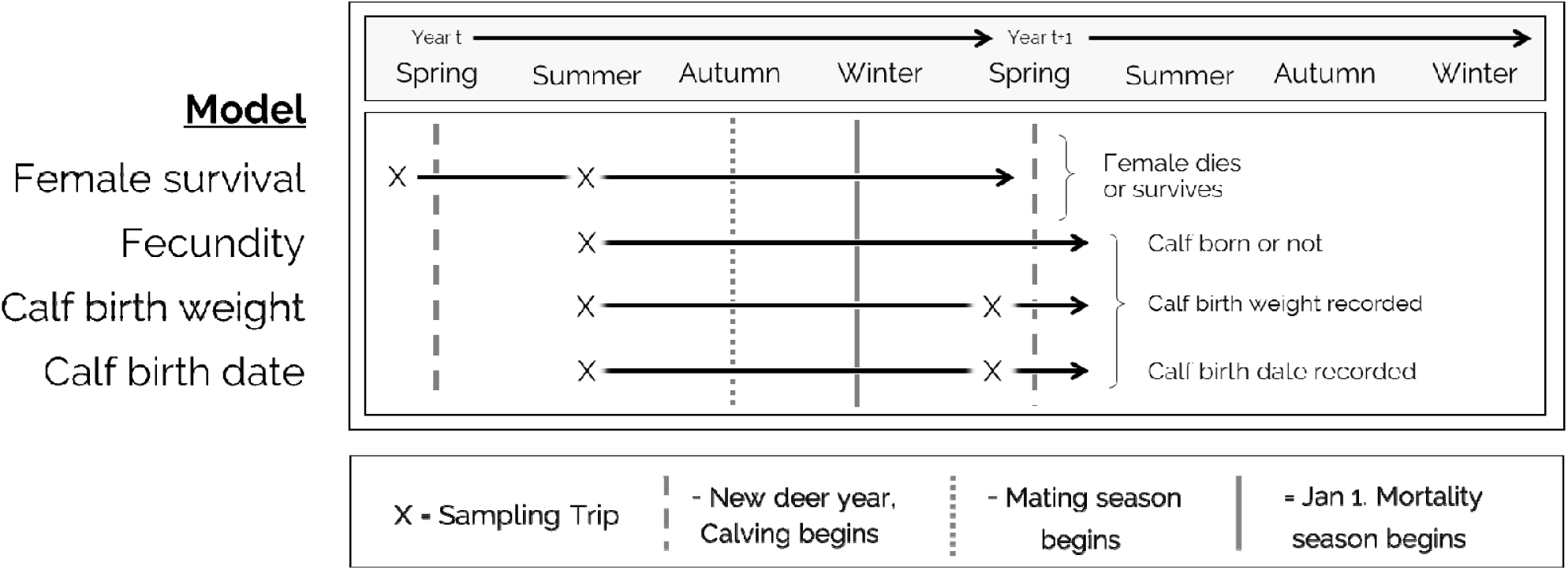
The four models in the context of the red deer reproductive cycle and sampling regime, over an example two-year period. Reproduction begins in spring and summer, at the start of each deer year, and one sampling trip was undertaken each summer (August), after the calving season had finished. Mating occurs in the autumn, and the mortality season begins in winter and lasts until early spring. A second sampling trip occurred each spring (April), after mating and mortality, but before the beginning of the subsequent calving season. The fitness variables investigated were quantified at the start of the subsequent deer year: if a female survived to May 1 the following year she was counted as 1 in the survival analysis, 0 if not, and the presence, weight, and birth date of her calf in the following spring were used as response variables in the remaining three models. The sampling trips included in each model were selected according to feasibility of causal links. For example, females become pregnant in the autumn, so we did not include the spring sampling season in the reproduction model as they would already be pregnant at this point, making it unlikely that parasite counts in April have a direct effect on their probability of having a calf 1-2 months later.

During early spring (April) and late summer (August), either side of the calving season, we conducted two-week field trips to collect faecal samples from the deer noninvasively. Sampling was undertaken across five trips 2016-2018, with 701 faecal samples collected in total; see Table 1 for details of datasets. We watched known individuals for defaecation, marked the spot where the droppings landed, and then collected them while minimising disturbance to the deer, generally within an hour. In the evenings, samples were processed and put into Ziploc bags for storage (Albery et al 2018). A subsample was extracted by centrifugation and kept frozen for faecal antibody analysis (Watt *et al.*, 2016; Albery *et al.*, 2020). Another subsample was kept as anaerobic as possible in a Ziploc bag at 4°C to avoid egg hatching.

**Table 1:** Descriptions of path analyses and the datasets used.

| Model Set | Fitness Measure | Definition | Dataset | Samples | Individuals |
| --- | --- | --- | --- | --- | --- |
| 1 | Survival | Female survival the following winter (0/1) | All females (Spring year t-1 and Summer year t) | 485 | 134 |
| 2 | Fecundity | Female reproduction the following deer year (0/1) | All females (Summer) | 223 | 107 |
| 3 | Calf birth weight | Calf weight the following deer year (Kg) | Females that reproduced the following May-June (Summer year t and Spring year t) | 300 | 94 |
| 4 | Parturition date | Date of parturition the following deer year (Days from 1<sup>st</sup> January) |  | 336 | 106 |

In the lab, faecal samples were counted for strongyle nematodes eggs using a salt flotation-centrifugation faecal egg count (FEC), accurate to 1 egg per gram (EPG) within 3 weeks of collection. Strongyles are ubiquitous ruminant parasites that are present at high prevalence in this population and which increase in parasitism in lactating individuals (Albery *et al.*, 2020). Previous studies in this population have also examined the helminths *Elaphostrongylus cervi* and *Fasciola hepatica* (Albery *et al.*, 2018, 2020). We chose to examine strongyles but not *E. cervi* or *F. hepatica* for several reasons: we did not want to add too many links to the analysis for reasons of interpretability; strongyles are most expected to have strong fitness costs (Hoberg *et al.*, 2001) and exhibited the most profound reproductive tradeoff in terms of significance and magnitude (Albery *et al.*, 2020); we did not expect *E. cervi* to have strong fitness effects (Irvine *et al.*, 2006); and *F. hepatica* is present at relatively low prevalence in adult females, preventing it from being fitted easily as an explanatory variable (Albery *et al.*, 2018). We also carried out antibody detection ELISAs designed to quantify mucosal IgA in sheep (Watt *et al.*, 2016; Albery *et al.*, 2020). This protocol quantifies both total IgA levels as a measure of general immune investment, and anti-*Teladorsagia circumcincta* IgA levels (anti-Tc IgA) as a specific anti-strongyle measure. *T. circumcincta* is primarily a sheep strongyle, but the anti-Tc IgA assay shows high cross-reactivity with a range of strongyle nematodes including the mouse helminth *Heligmosomoides polygyrus* (Froy *et al.*, 2019). The deer are infected with a selection of strongyle nematodes (Irvine *et al.*, 2006), including *Teladorsagia circumcincta* (unpublished data); thus, anti-Tc IgA is used to approximate anti-strongyle immune responses in the deer (Albery *et al.*, 2020). To control for collection factors which introduce confounding variation in antibody levels we used the residuals from a model including extraction session, time to freezing, and collection day, as in previous studies (Albery *et al.*, 2019, 2020). We also assayed faecal samples collected in November (Albery *et al.*, 2018). However, females exhibited very low strongyle prevalence in the autumn compared with spring and summer, preventing our FEC data from approximating normality and providing little variation to test when fitted as an explanatory variable. Hence, autumn data were excluded from our analyses.

### Statistical analysis

To investigate links among our variables we used path analysis using the D-sep method, in which a set of linear models are fitted to the data, with some variables appearing as both response and explanatory variables (Shipley, 2009). Combining the linear models in this way allows identification of potential causal links and mediating variables.

We created four Directed Acyclic Graphs (DAGs), each examining a different fitness-related trait measured in the year following measurement of parasite burden (see Figure 2). These measures included two direct fitness measures: the female’s overwinter survival (0/1, where 1=survived to May 1 the following year) and fecundity the following year (0/1, where 1=gave birth in the following calving season). We also examined two fitness-associated maternal traits: the birth weight of a female’s calf the following year (continuous, Gaussian distributed, based on a regression of capture weight on capture age in days) and parturition date the following year (continuous, Gaussian distributed, based on Julian date that year).

Our analyses used three immune and parasite measures, which included: Total IgA level; Anti-Tc IgA level; Strongyle count per gram of faeces (continuous, log(count+1)-transformed to approximate normality). We included two mutually exclusive binary reproductive categories representing the reproductive cost paid that year (Clutton-Brock *et al.*, 1989): Gestation (gave birth to a calf which died before 1^st^ October that year) and Gestation + Lactation (gave birth to a calf which survived to 1^st^ October; hereafter referred to as simply “Lactation”, as all individuals that lactated must have also undergone the cost of gestation). We also included variables to control for annual, seasonal, and age-related variation: Year (categorical, with three levels: 2015, 2016, 2017); Season (two levels: Summer, Spring); and Age (continuous, in years).

Each of the four DAGs was composed of four similar models, fitted using the INLA package (Rue and Martino, 2009) in R version 3.5 (R Core Team, 2018). All measures included female identity as a random effect to control for pseudoreplication. First, we ran a set of three “input models”, where the response variable was an antibody or parasite measure. The aim of these models was to quantify the association between reproduction and the immune/parasite measures, and to quantify links between these measures themselves.

The models were specified as follows for each of our analyses, with immune/parasite measures in bold and reproductive traits in italics. Variables in brackets were included in the models, but are not displayed in the DAGs for clarity.

1. **Total IgA** ∼ *Gestation* + *Lactation* (+ Age + Season + Year)
2. **Anti-Tc IgA** ∼ **Total IgA** + *Gestation* + *Lactation* (+ Age + Season + Year)
3. **Strongyles** ∼ **Anti-Tc IgA** + *Gestation* + *Lactation* (+ Age + Season + Year)
4. Fitness-related trait ∼ **Strongyles** + **Anti-Tc IgA** + **Total IgA** + *Gestation* + *Lactation* (+ Age + Year)

Combining these two model sets allowed comparison of the significance and magnitude of different traits’ costs for fitness in the following year (Figure 1). Combining the estimates from models 1-3 with the estimates from model 4 allows calculation of the direct and indirect (parasite- or immune-mediated) effects of lactation and gestation on subsequent fitness traits (Figure 1). As an example, we compared the magnitude and credibility intervals of direct lactation effects (effect of lactation in the fitness model [model 4]) with indirect effects (lactation effects on strongyle count [model 3] multiplied by the effects of strongyle count on fitness [model 4]). We took 1000 posterior draws from each of the lactation-strongyle link and the strongyle-fitness link and multiplied them together, and then derived the 95% credibility intervals for this link. We compared these estimates with those for the direct lactation-fitness link to investigate whether effects of lactation were likely to act independently and/or through strongyle count. The models, fitness measures, and datasets used in each analysis are described in Table 1.

## Results

Path analyses consistently revealed strong positive associations between lactation and parasite count, and negative associations between parasite counts and subsequent fitness in terms of all four traits (Figures 3-5). In contrast, estimates for lactation’s direct association with subsequent fitness overlapped with zero for all response variables except parturition date, supporting parasite-mediated reproductive costs for fitness (Figures 3-5). Below, for each of the four fitness-related response variables, we describe the magnitude of the direct association of parasitism with fitness, the direct association of lactation with fitness, and lactation’s association with parasitism multiplied by parasitism’s association with fitness. The latter gives an estimate of the indirect effect of lactation on the fitness-related trait acting through strongyle count. For effect sizes we give the mean and 95% credibility intervals (CI). 1 log(EPG+1) increase corresponds to a ∼3x increase in strongyle count. Full model effect sizes are displayed in the supplementary information (Figure SI1; Table SI1).

**Figure 3:**
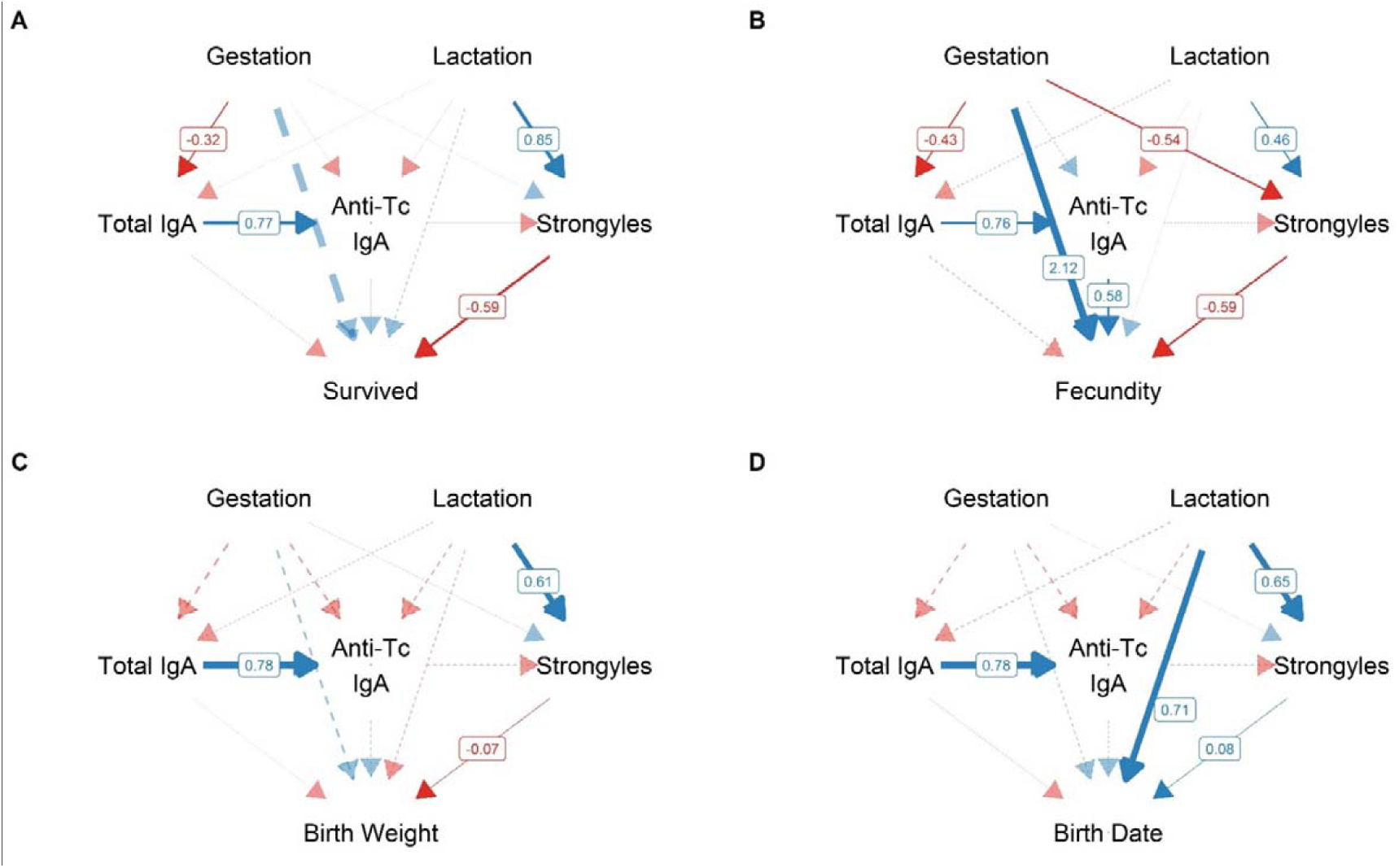
Directed Acyclic Graphs (DAGs). Results are displayed for all four investigated fitness response variables: overwinter survival (A); fecundity (B); subsequent calf birth weight (C); subsequent parturition date (D). Link colour depends on the direction of the effect (blue=positive, red=negative); link width indicates the magnitude of the effect; and only solid, opaque links are significant (estimates did not overlap with zero). Labels denote the link-scale effect sizes (slopes) for the significant effects, derived from GLMMs (full model effects displayed in SI Figure 1).

Parasite count had a strong association with subsequent survival probability despite high survival rates in the population (Figure 3A,4A). Females with the lowest counts (0 EPG, 10% of samples) had a survival probability of ∼100%, while those with the highest (>25EPG, 7% of samples) had a survival probability of <90% (Figure 4A). Lactation was associated with increased strongyle count (+0.85, CI 0.64, 0.99), so that a substantial cost of lactation for survival acted through parasitism (Figure 5). Although this effect was highly significant on the link (logistic) scale (Figure SI1, Table SI1), given the high survival rates in the population, at the mean EPG value this lactation-associated increased strongyle parasitism would correspond to only a ∼2% decrease in survival probability. In contrast, estimates for the direct effect of lactation on survival overlapped widely with zero, and the point estimate was greater than zero, implying that individuals that lactate were slightly more likely to survive when the effects of parasitism were accounted for (Figures 3A, 5).

**Figure 4:**
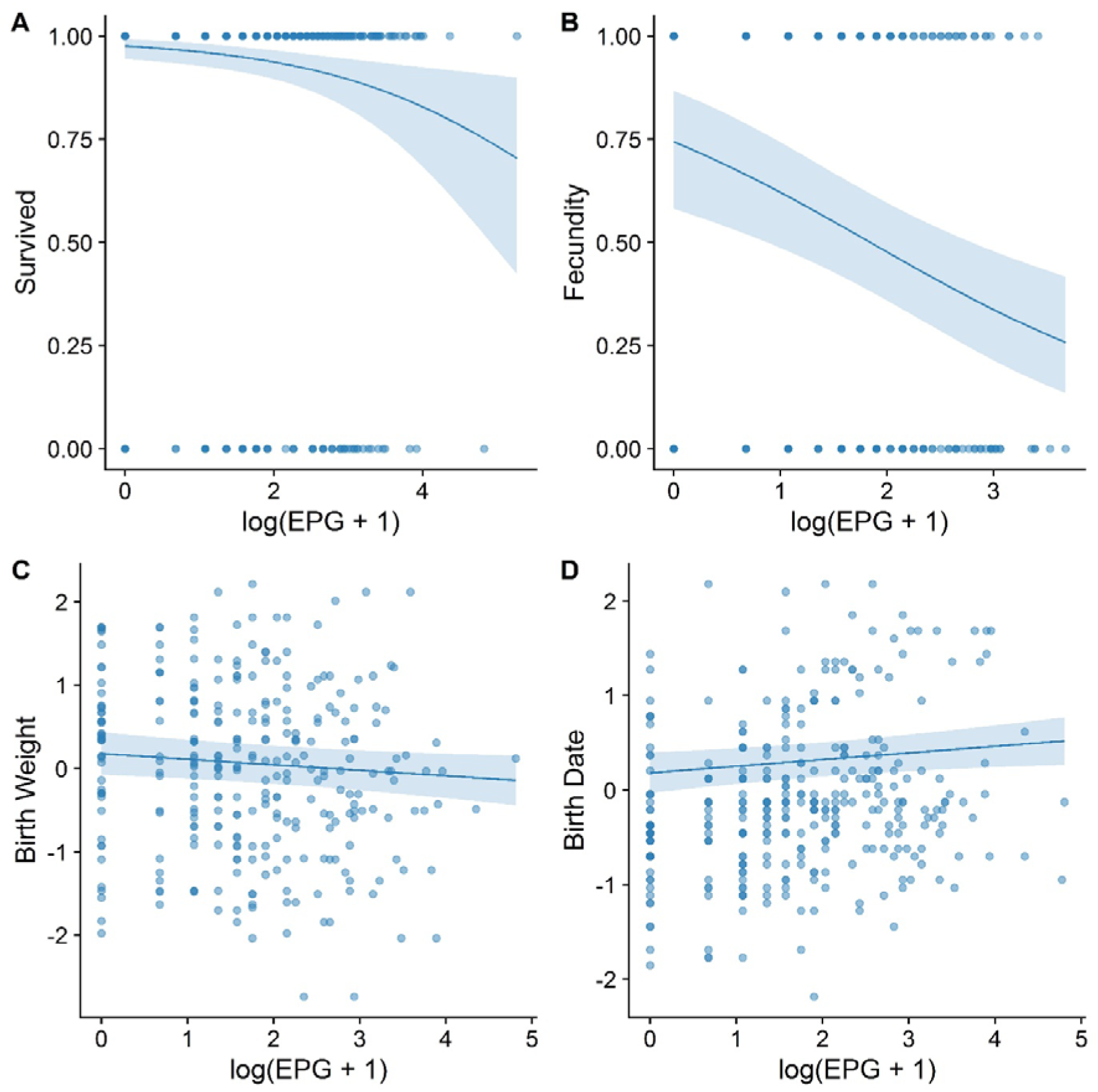
Relationships between strongyle parasite count and fitness measures. Results are displayed for all four investigated fitness response variables: overwinter survival (A); fecundity (B); subsequent calf birth weight (C); subsequent parturition date (D). The lines denote the fitted slope of parasitism on the response variable, with 95% credibility intervals. Credibility intervals did not overlap with zero for any of the four relationships. Strongyle count was log(x+1)-transformed for analysis and plotting.

**Figure 5:**
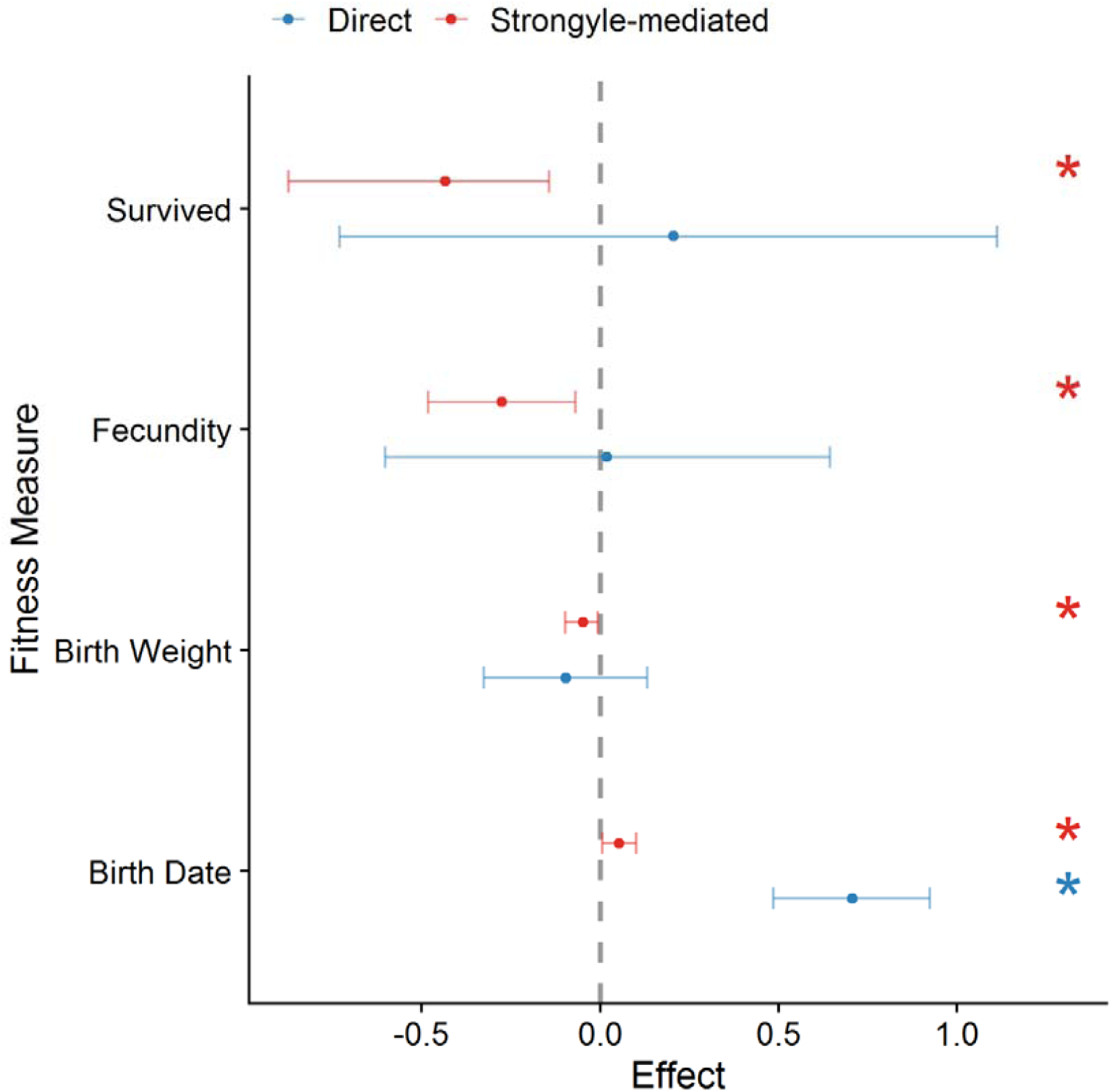
Comparison of direct and indirect (parasite-mediated) effects of lactation on fitness-related traits on the link scale (logistic for survival and reproduction; Gaussian for birth weight and birth date). Points represent mean effect estimates derived from the model posterior distributions; blue corresponds to direct effects, and red corresponds to indirect effects. Parameters with asterisks were significant: i.e., their credibility intervals did not overlap with zero.

Strongyles’ association with subsequent fecundity had a similar effect size to its association with overwinter survival (Figures 3B, 4B, 5; Table SI1; Figure SI1). An increase of 1 log(EPG+1) was associated with a decrease of ∼15% probability of reproducing. 0 EPG (17% of samples) corresponded to a ∼77% chance of reproducing the following year, and those with >20 EPG (6% of samples) had a reproduction probability of <36% (Figure 4B). The direct effect of lactation on subsequent fecundity was negligible and had very large credibility intervals, as with survival (Figure 5). In addition to the association with parasite count, individuals with higher levels of anti-Tc IgA were more likely to reproduce the following year (Figure 3B; Figure 6). An increase of 1 standard deviation of anti-Tc IgA levels corresponded to an increase of ∼10% in the probability of reproducing. Individuals with the lowest anti-Tc IgA levels (less than −1 SD units) had a reproduction probability of <50%, compared to >75% for those with the highest levels (>1 SD units; Figure 6). Finally, individuals that paid the cost of gestation were much more likely to reproduce the following year, independently of the effects of antibodies and parasites (Figure 3B).

**Figure 6:**
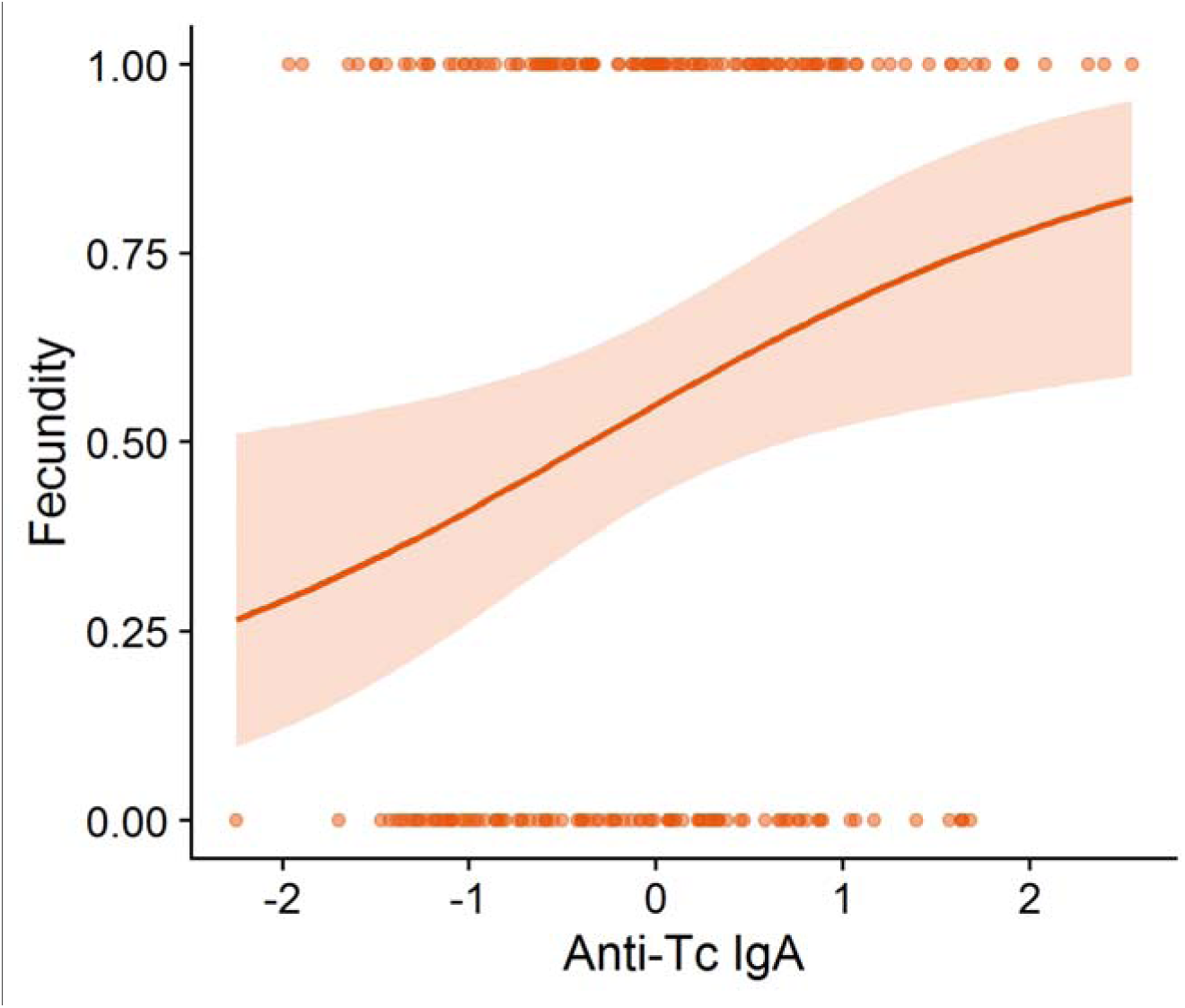
Higher anti-*Teladorsagia circumcincta* IgA was associated with an increased probability of reproducing the following year. Anti-Tc IgA was cube root-transformed and calculated from the residuals of a linear model including collection variables, and was then scaled to have a mean of 0 and a standard deviation of 1. The line represents the output of the reproduction probability model which includes lactation and strongyles as explanatory factors.

Calving traits exhibited weaker associations with parasitism than did survival and reproduction, although the results still implied an indirect cost of lactation acting through strongyle count (Figures 3-5). The DAG for calf birth weight was similar to that for survival (Figure 3C). An increase of 1 log(EPG+1) corresponded to a slight decrease in calf birth weight the following year (0.07 SD units, or about 86g; Figure 4C). Females with the highest strongyle intensities (>25 EPG) gave birth to calves which were ∼400g lighter than those with the lowest intensities (0 EPG), or around 6.24 kg compared to 6.65 kg. As with survival, there was poor support for a direct association between lactation and birth weight (Figure 5). The estimates for this direct effect were close to zero, and credibility intervals overlapped substantially with zero (Figure 5). Lactation’s positive effect on strongyle count once again resulted in a significant negative indirect effect of lactation on subsequent calf birth weight acting through strongyles, but the estimates were very small and nearly overlapped with zero (−0.0438kg, CI −0.111.6, −0.005.6).

In contrast to all other metrics we investigated, there was support for a positive and direct effect of lactation on parturition date the following year: that is, females whose calf survived until the winter were likely to calve later in the following year (∼8.5 days later, CI: 5.9, 11.2; Figure 3D,5), regardless of parasite count. There was a much weaker association between strongyle count and parturition date: an increase of 1 log(EPG+1) produced a delay in calving of ∼0.93 days (CI: 0.12, 1.75; Figure 3D,4D,5). Lactation resulted in an increase of 0.7 log(EPG+1). Combining this estimate with the effect of parasitism on birth date gives an estimate for an indirect effect of lactation acting on birth date totalling 0.58 days’ delay (CI 0.06, 1.31; Figure 5). Parturition date was thus the only metric examined here for which lactation’s direct effect was larger than its indirect effect acting through strongyle count (Figures 3-5).

There was a strong positive association between total IgA and anti-Tc IgA, as expected given our previous findings (Albery *et al.*, 2020; Table SI1, Figure SI1). However, lactation had no significant effect on anti-Tc IgA in our DAGs (Figure 3, Table SI1, Figure SI1).

## Discussion

This study provides observational evidence for strong parasite-dependent mediation of multiple life history tradeoffs in a large wild mammal. Lactation was associated with higher parasite intensities which translated to reduced fecundity and survival probability in the subsequent year. Among individuals that did reproduce the following year, those with high strongyle counts gave birth slightly later in the year and to slightly smaller calves. These findings represent the second evidence for such mediation of reproduction-survival tradeoffs in a wild mammal (Leivesley *et al.*, 2019), and new evidence of parasites mediating reproductive tradeoffs with subsequent reproductive traits. It is likely that much of the fitness reduction associated with lactation in the Rum red deer population (Clutton-Brock *et al.*, 1989; Froy *et al.*, 2016) is caused by strongyle parasites, or that strongyle count closely corresponds to latent condition variables that are responsible for mediating fitness. This finding supports parasites’ role as an important mediating factor in this system.

Lactation’s negative association with fitness acted largely through strongyle count for all fitness metrics except parturition date. This may represent a parasite-mediated cost, where pathology and resource allocation associated with increased parasitism are the primary cause of increased overwinter mortality and reduced subsequent fecundity in lactating individuals (Clutton-Brock *et al.*, 1989). Allocation of resources to lactation and associated physiological changes likely reduces resources available for resistance and damage repair mechanisms, rendering lactating females more susceptible to strongyles (Sheldon and Verhulst, 1996; Speakman, 2008), while also increasing their exposure through heightened forage intake (Albery *et al.*, 2020). High parasite counts in lactating females may cause gut pathology, interfering with nutrient absorption and thereby exacerbating the nutritional scarcity of the winter period, leading to overwinter mortality (Gulland, 1992; Pedersen and Greives, 2008; Maublanc *et al.*, 2009), as well as reducing females’ ability to achieve the body condition necessary to conceive and carry a calf to term (Albon *et al.*, 1986). This reduction in body condition could likewise cause females to give birth later in the year and to a calf that is smaller. There are two time points at which strongyles may reduce fecundity: first, parasites may impact females in the resource-abundant summer and early autumn, preventing them from conceiving in the autumn mating season. In this case, strongyle-associated pathology may occur somewhat independently of overwinter nutritional scarcity. Alternatively, strongyles may cause females to lose their pregnancies over winter. This possibility may be tested in the future by investigating whether more highly parasitised females are less likely to be observed mating (demonstrating reduced conception rates), or only less likely to give birth.

Lactation exerts a substantial resource cost that results in reduced condition; therefore, it is also possible that we observed a negative relationship between parasitism and fitness because both were determined by condition, rather than because parasites were causally responsible for reducing fitness. Strongyle counts are associated with decreased body condition in shot individuals in this population, supporting this possibility (Irvine *et al.*, 2006). Similarly, an important role of condition is supported by our observation that higher anti-Tc IgA levels were associated with increased fecundity the following year, independently of any associations between 1) strongyles and fecundity and 2) anti-Tc IgA and strongyles themselves (Figure 6). It is highly likely that anti-Tc IgA is well-correlated with an unmeasured component of individual quality such as fat content (Demas *et al.*, 2003) which is linked to fitness both in the deer (Albon *et al.*, 1986) and in other systems (Milenkaya *et al.*, 2015). This possibility reflects the confounding effects of individual quality in observational studies of tradeoffs (van Noordwijk and de Jong, 1986). We were unfortunately unable to replicate previous findings of lactation costs for mucosal antibodies (Albery *et al.*, 2020), likely due to extremely reduced sample sizes (485 samples and fewer here compared to 837 samples previously), and so we were unable to link the anti-Tc IgA-fitness association as part of a reproduction-fitness tradeoff. Another potential fitness-mediating factor is body weight, which is often used to control for condition-driven versus parasite-driven fitness effects; however, analyses in Soay sheep often show that strongyles-fitness associations occur independent of, or in addition to body weight (e.g. Sparks *et al.*, 2018; Froy *et al.*, 2019; Leivesley *et al.*, 2019). In addition, although condition-parasitism relationships are well-documented, negative effects are far from ubiquitous and their slopes are relatively shallow on average (Sánchez *et al.*, 2018). Whether or not strongyles are the effectors, our findings nevertheless support the use of these parasites as a proxy for an individual’s health and as predictors of its subsequent fitness.

Strongyles will have a strong mediating effect on population dynamics, for two principal reasons: first, by reducing both survival and fecundity simultaneously, and second, by exhibiting different relationships with past and subsequent reproduction. As such, it stands to reason that their impact will prevent too many females from reproducing in a single year, potentially stabilising population fluctuations. Further years of data will reveal how parasite abundances relate to the population dynamics of the deer, and particularly whether inter-annual variation in strongyle numbers can explain population size (Wilson *et al.*, 2004). At higher population densities the deer exhibit delayed maturity and reduced fecundity (Albon *et al.*, 1983); the lactation-strongyle-fecundity tradeoff offers a potential mechanism behind this fecundity reduction, particularly as parasitism should worsen at higher densities (Altizer *et al.*, 2003; Wilson *et al.*, 2004). Local population density also influences fitness in this population (Coulson *et al.*, 1997) and parasitism demonstrates fine-scale spatial variation (Albery *et al.*, 2019), so this life history mediation could likewise occur at relatively fine spatial scales. A similar study in Soay sheep demonstrated that strongyles mediate a reproduction-survival tradeoff, but without examining similar reproduction-fecundity tradeoffs, partly because most sheep do not take years off between reproduction events (Leivesley *et al.*, 2019). The fecundity reduction seen in the deer and the strength of these parasite-mediated tradeoffs potentially contribute to the population’s relatively weak population cycles, particularly compared to the strong oscillatory population dynamics of the Soay sheep (Clutton-Brock and Pemberton, 2004). As such, parasite-dependent life history mediation may be an important contributing factor determining the strength of oscillatory population dynamics.

Finally, having uncovered costs of parasitism in adult females, it would be interesting to investigate whether other age and sex categories experience similar fitness effects: e.g., do more highly parasitised males sire fewer calves, and are more highly parasitised calves less likely to survive to maturity? Do maternal costs transfer to their calves, providing another potentially important mediating mechanism (Martin and Festa-Bianchet, 2010)? Future studies in this population could elaborate on these findings by investigating how maternal and calf parasitism correlate and correspond to maternal and calf fitness, quantifying transgenerational immunity-parasitism-fitness correlations: a topic that is largely understudied and likely influences ecological and epidemiological dynamics considerably (Roth *et al.*, 2018).

## Supplementary Information

**Figure SI1:**
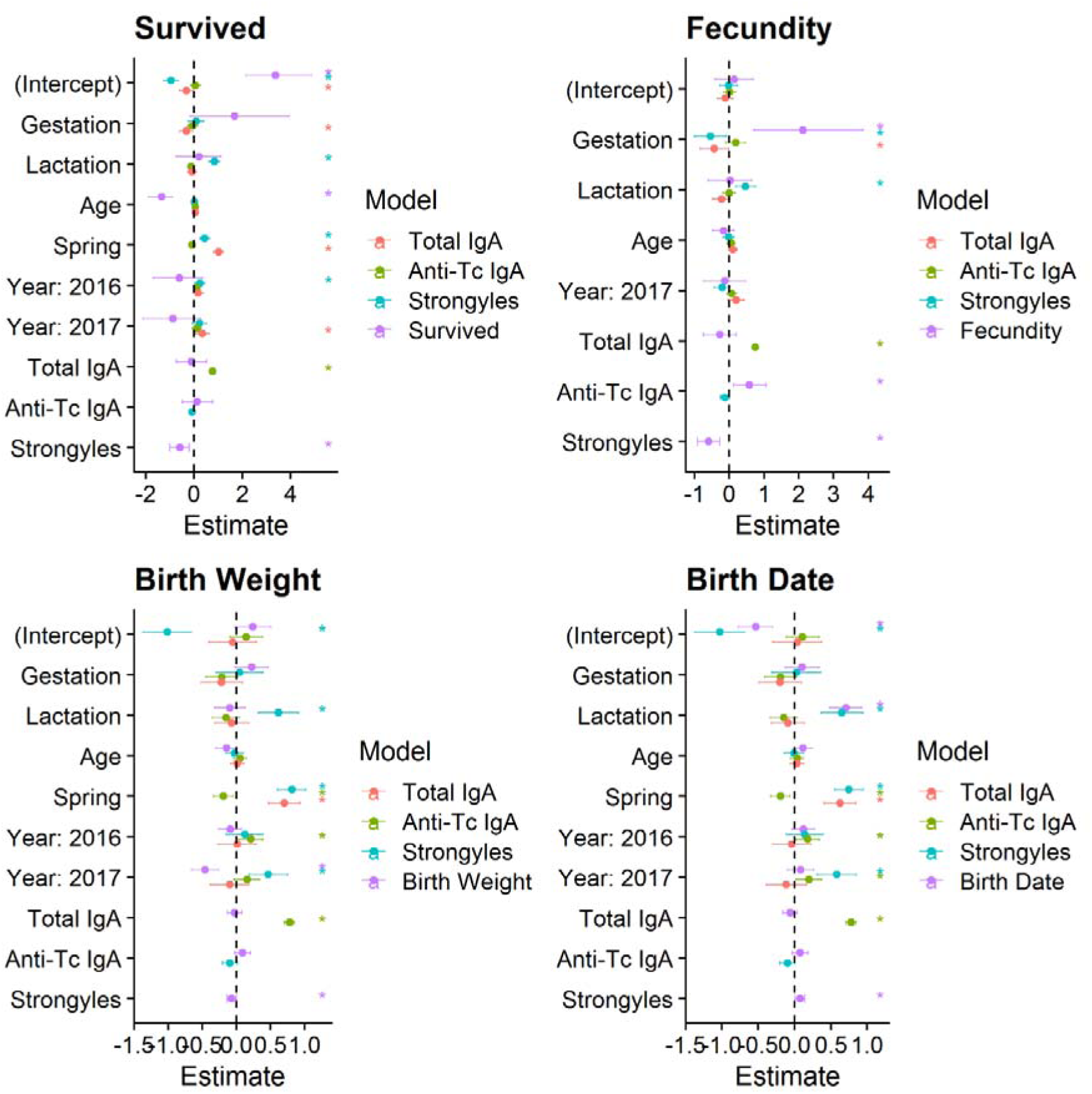
Model outputs for the component GLMMs of all four path analyses. Colours correspond to different response variables; variables on the y axis correspond to explanatory variables in each component model. Points denote mean estimates from the posterior distribution of effect sizes from the component INLA model. Error bars denote 95% credibility intervals for the distribution. Where these intervals did not overlap with 0, the effect was taken to be significant. These effects are marked with an asterisk. Notably, lactation was positively associated with strongyles (blue estimates) and strongyles were negatively associated with all fitness-related traits (purple estimates).

**Table SI1:**
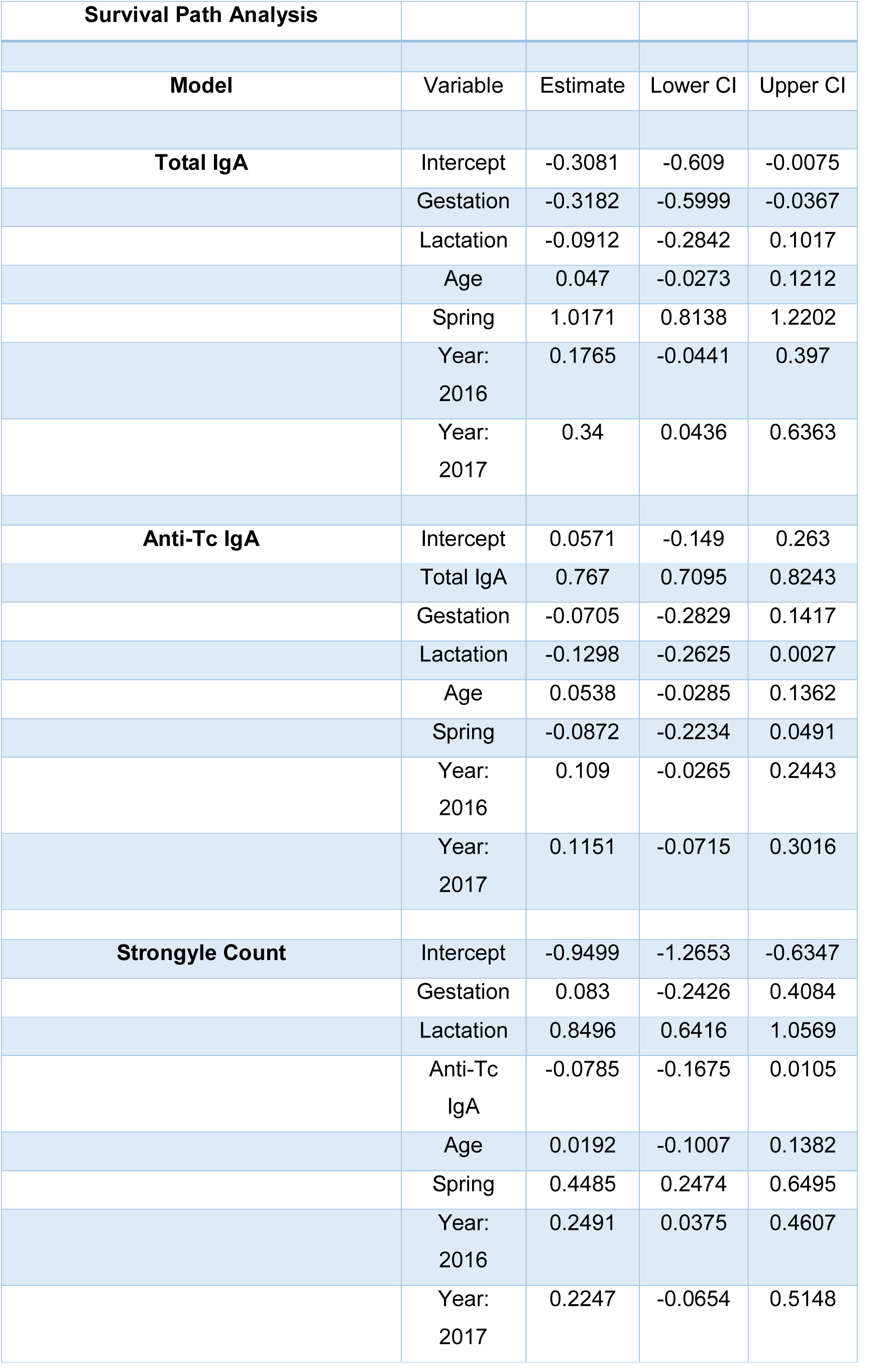
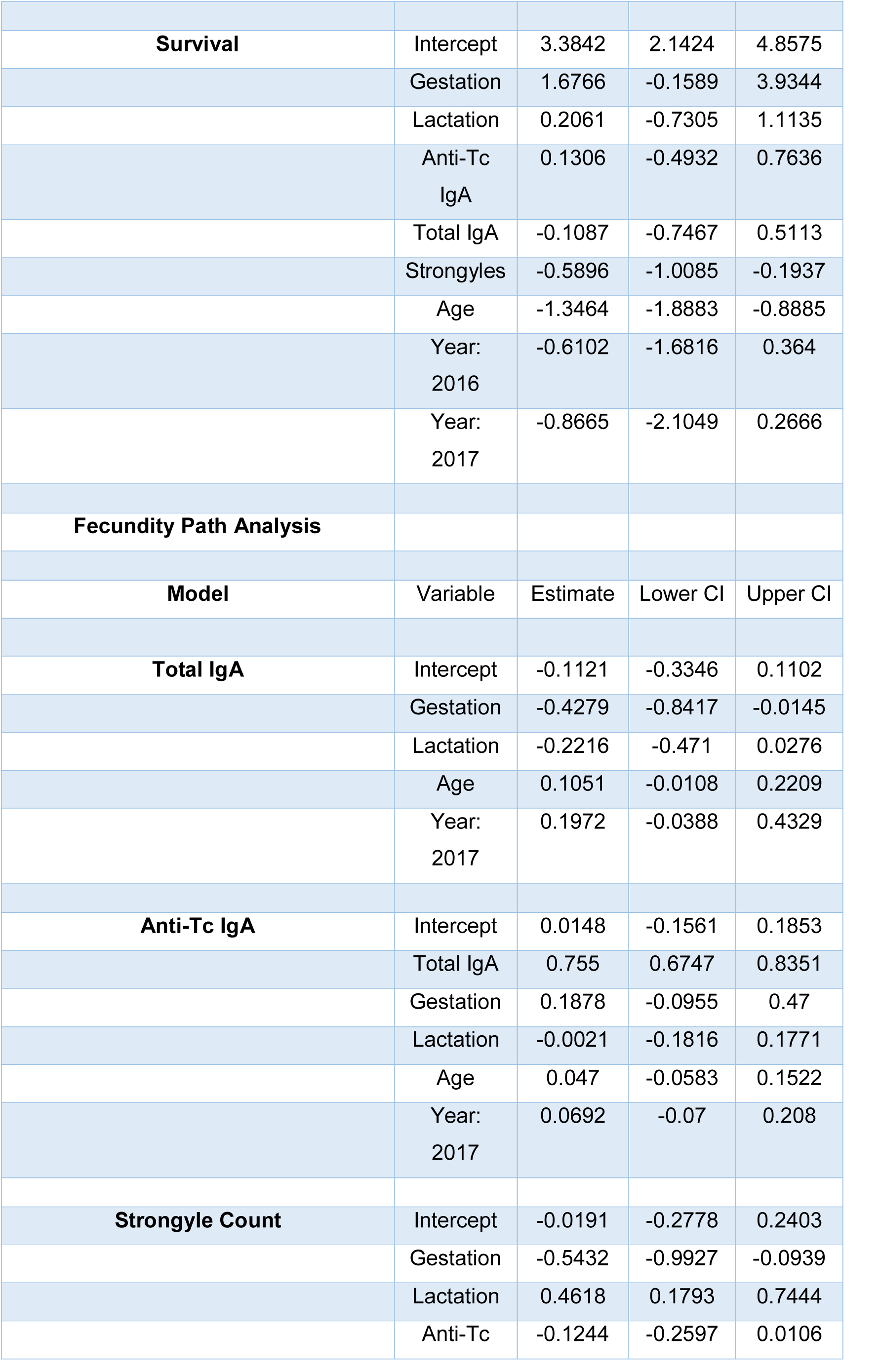
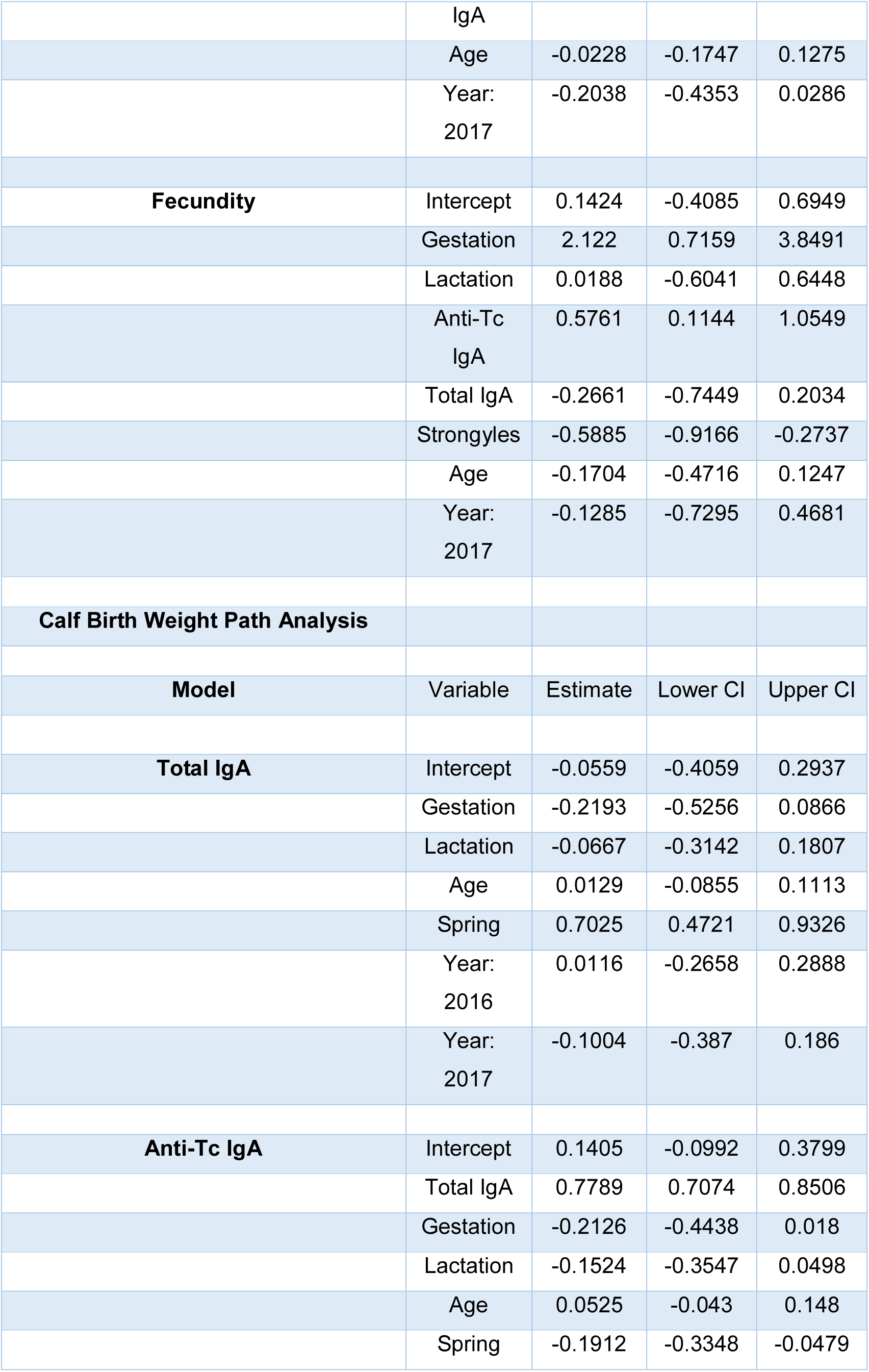
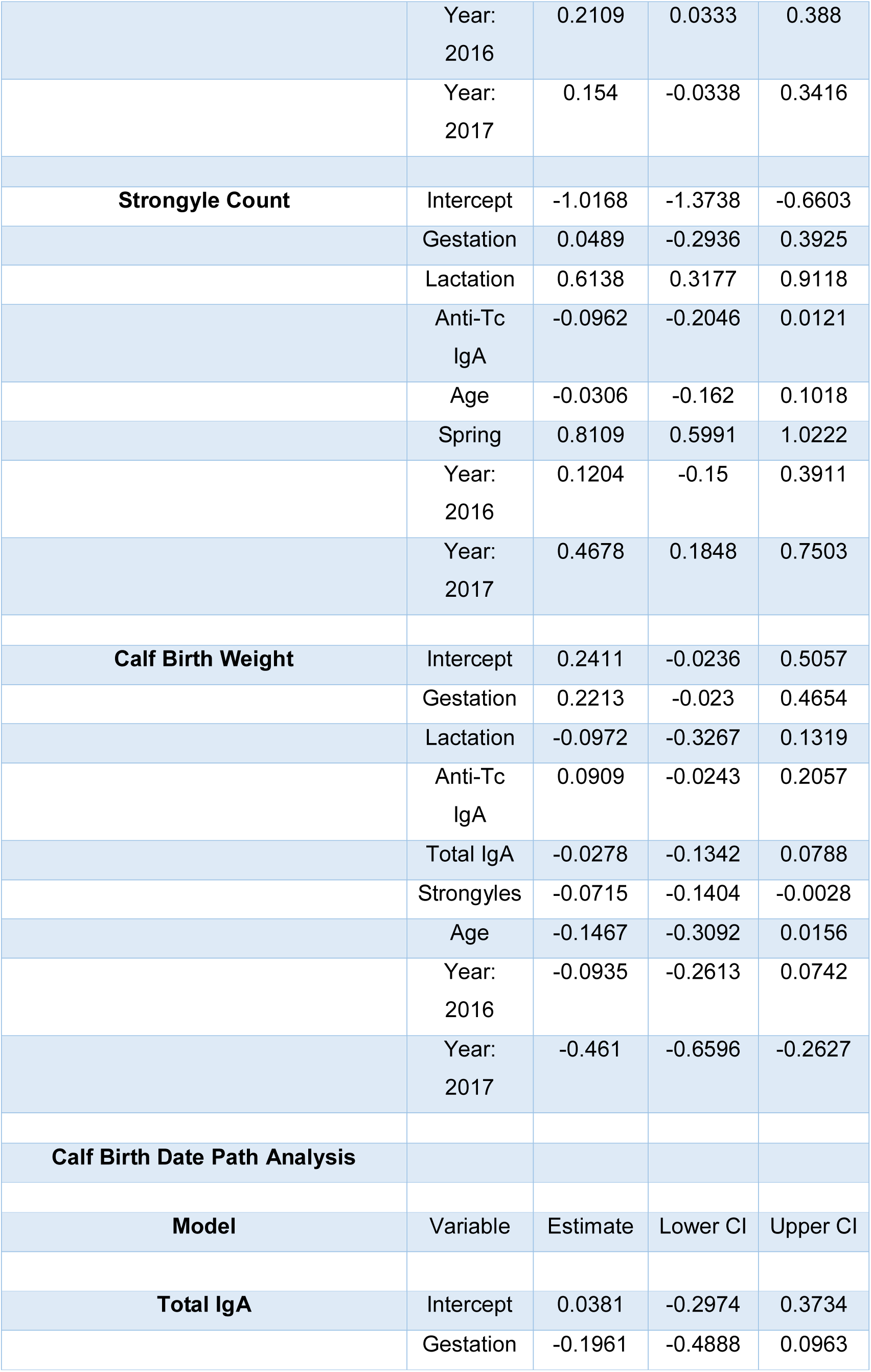
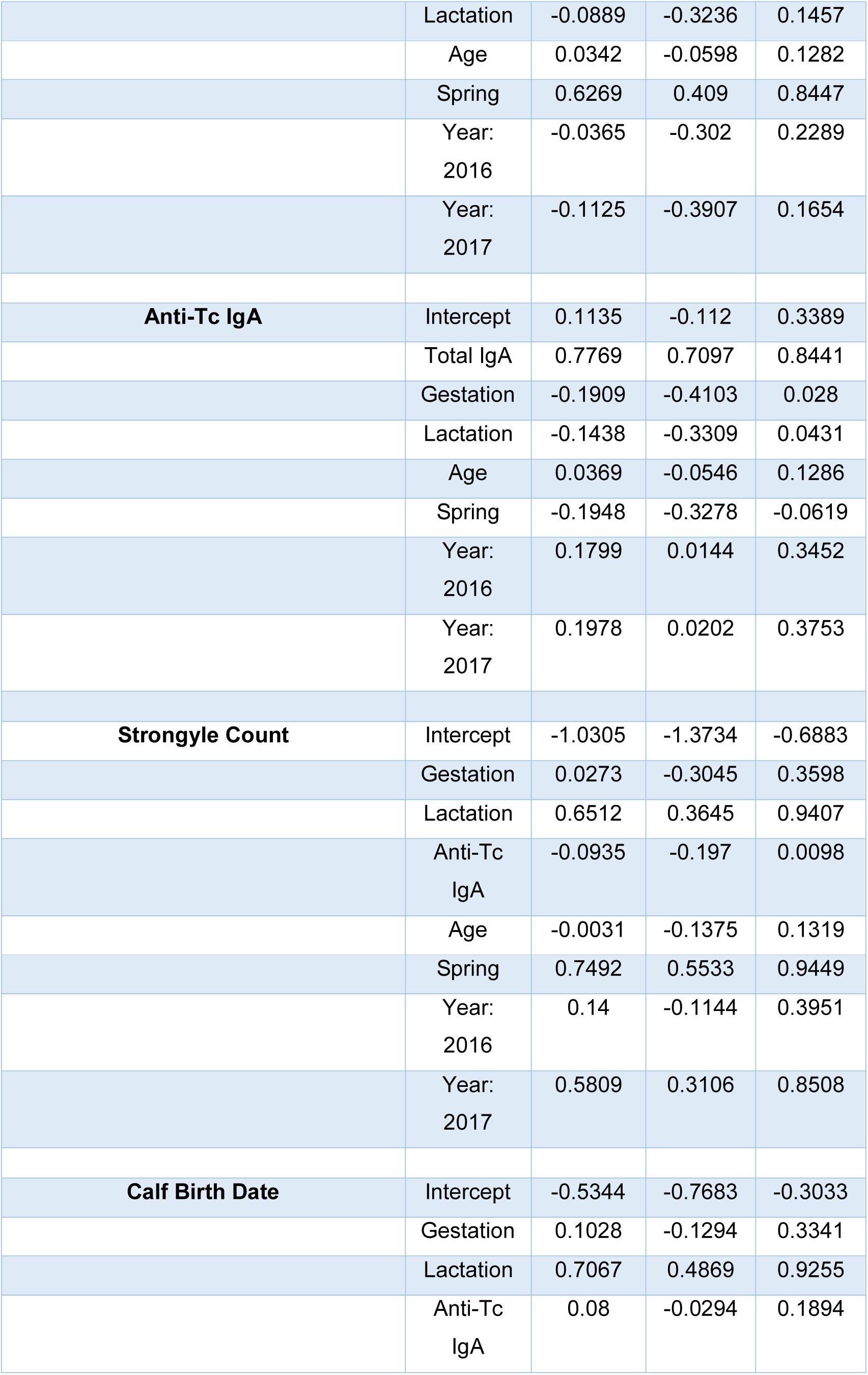
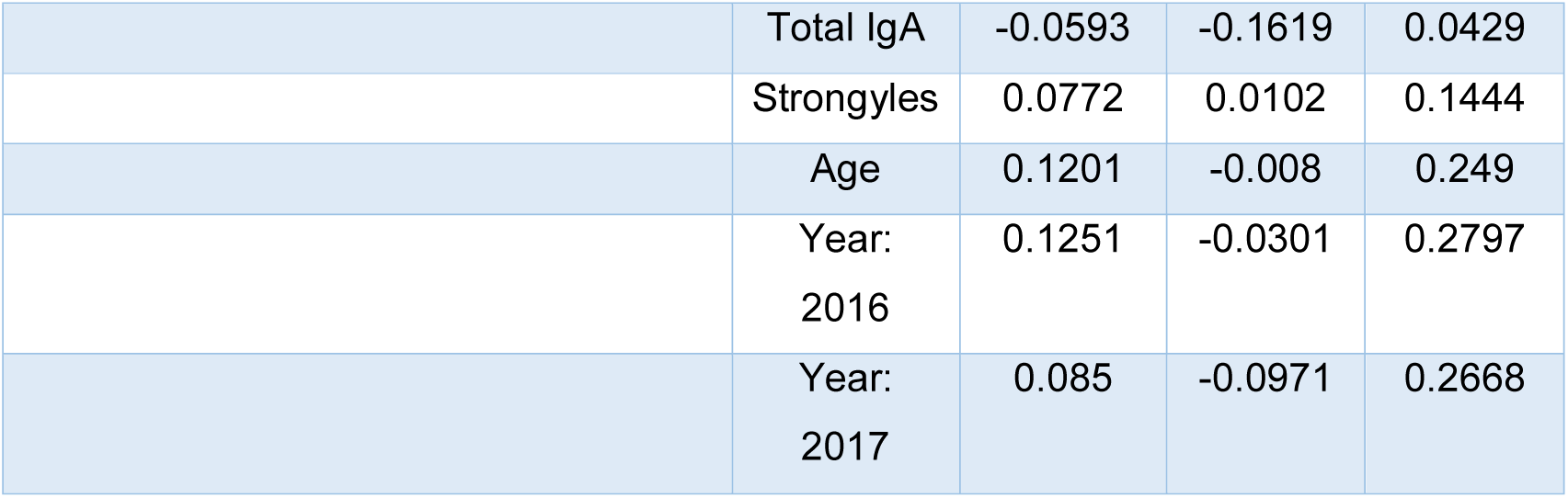
effect sizes and 95% credibility intervals for the component GLMMs of each path analysis

